# Continuous assembly required: perpetual species turnover in two trophic level ecosystems

**DOI:** 10.1101/2023.03.08.531662

**Authors:** Jurg W. Spaak, Peter B. Adler, Stephen P. Ellner

**Author notes:** Email addresses. Statement of authorship*: J.W.S. wrote the computer code. J.W.S. wrote the first draft, all authors discussed content and contributed to revisions. Open Research statement*: We created new computer code for the simulations of this manuscript. The computer code is publicly available at https://zenodo.org/record/7544362#.Y8af2HaxND8 with DOI 10.5281/zenodo.7544362 Upon acceptance we will store the computer code on Zenodo. We did not use any new empirical data in this manuscript.

## Abstract

Community assembly is often treated as deterministic, converging on one or at most a few possible stable endpoints. However, in nature we typically observe continuous change in community composition, which is often ascribed to environmental change. But continuous changes in community composition can also arise in deterministic, time-invariant community models, especially food web models. Our goal was to determine why some models produce continuous assembly and others do not. We investigated a simple two trophic-level community model to show that continuous assembly is driven by the relative niche width of the trophic levels. If predators have a larger niche width than prey, community assembly converges to a stable equilibrium. Conversely, if predators have a smaller niche width than prey, then community composition never stabilizes. Evidence that food webs need not reach a stable equilibrium has important implications, as many ecological theories of community ecology based on equilibria may be difficult to apply to such food webs.

## Introduction

Understanding how species assemble into communities is a central issue in community ecology (Fukami, 2015; Song *et al*., 2021; Serván & Allesina, 2021). Community assembly is typically modeled as a sequence of invasions of species from a regional species pool into a local patch, where the success of each invasion may depend on both the environmental conditions as well as the local community itself (HilleRis-Lambers *et al*., 2012; Barbier *et al*., 2021).

How we view community structure affects how we interpret community assembly (Tilman, 2004). A view based on niche theory typically implies a deterministic community assembly process, where composition converges on the community best adapted to the environment (MacArthur, 1970; Tilman *et al*., 1982; Cressman *et al*., 2017; Kremer & Klausmeier, 2017). For example, with competition for a single limiting resource, the species with the lowest resource requirement R* replaces all its competitors with higher resource requirements (Tilman *et al*., 1982; Tilman, 2004). Similar rules allow us to predict community assembly under competition for two resources (Tilman *et al*., 1982), with predators (Holt & Lawton, 1994) or with mutualists (Johnson & Bronstein, 2019). In these models, resource competition leads deterministically to a single community in which every available niche is occupied (Tilman, 2004; Cressman *et al*., 2017), independent of the assembly processes.

Communities with priority effects, or other historical contingencies, are no exception: they also converge towards a predictable outcome of community assembly (Fukami, 2015; Serván & Allesina, 2021). However, in these cases the outcome can depend on the starting point and potentially on the community assembly process itself. Understanding when and how the sequence of community assembly affects the final community may not be simple (Fukami *et al*., 2016; Vannette & Fukami, 2014; Song *et al*., 2021; Huisman & Weissing, 2001; Barbier *et al*., 2021), but we still expect assembly to converge on one of several possible stable, uninvadable communities (Mordecai, 2011; Ke & Letten, 2018; Song *et al*., 2021).

Conversely, we rarely observe stable community compositions in natural communities (Blowes *et al*., 2019; Dornelas *et al*., 2019; Hamm & Drossel, 2021). Rather natural communities appear to be in a continuous community assembly process. Often, we observe a set of permanent species, typically called core species, and a set of transient species, typically called satellite species (Nee *et al*., 1991). Typical explanations for these patterns include environmental change (Dornelas *et al*., 2019) or neutral or stochastic processes (Hubbell, 2005). We accepted this view until recently when we investigated a two trophic-level plankton community model with mechanistic species interactions (Spaak *et al*., 2022). In this model, community composition changed continuously over time, despite the lack of external environmental changes or any stochastic processes.

As demonstrated by our plankton community model, patterns of continuous community assembly can also arise from internal species interactions in food web models that are purely deterministic and time-invariant (Hamm & Drossel, 2021; Morton & Law, 1997; Steiner & Leibold, 2004). Such models capture many of the patterns observed in nature such as food-chain length, number of average links per species, species-area relationships and average persistence time (Williams & Martinez, 2000; Loeuille & Loreau, 2005). However, not all food-web models lead to a continuous assembly pattern (Loeuille & Loreau, 2005) and some lead to a continuous assembly pattern only for higher trophic levels (Allhoff *et al*., 2015). The drivers of continuous community assembly are understood in some simple phenomenological models (Bunin, 2017), but these models are based on randomly generated matrices of species interaction coefficients, which do not reflect natural communities (Eklöf *et al*., 2013; Li *et al*., 2022). Interaction strengths in food web models and real food webs are highly structured, so continuous assembly in food web models is a different phenomenon. For example, predation strength in many food web models is based on a Gaussian function of differences in body sizes, yet while some of these models lead to continuous turnover others do not. Currently, we do not know which of the underlying assumptions of food web models are responsible for continuous community assembly.

Understanding the properties that lead to continuous community assembly is important, as many of our ecological theories are based on assumptions of stable community composition and equilibrium dynamics. For example, modern coexistence theory is based on invasions into stable communities at equilibrium (Ellner *et al*., 2019; Spaak *et al*., 2021; Barabás *et al*., 2018), studies of biodiversity-ecosystem function typically measure both biodiversity and ecosystem function at equilibrium (Loreau & Hector, 2001; Loreau, 2010; Bannar-Martin *et al*., 2018), and ecosystem stability analysis is based on linear approximations around an equilibrium (May, 1972; Carpentier *et al*., 2021; Allesina & Tang, 2012, 2015).

Here we analyze simple community models with one or two trophic levels (Macarthur & Levins, 1967; MacArthur, 1970) to answer two questions about continuous invasion and extinction dynamics. 1. What are the necessary conditions for these dynamics to emerge? 2. Are there any constant properties within the disorder of continuous invasion and extinction?

## Methods

### Community model and assembly

We first observed continuous invasion and extinction in a mechanistic phytoplankton-zooplankton model (Spaak *et al*., 2022). However, here we focus on a simpler two-trophic Lotka-Volterra community model because it offers greater generality and less complexity. The Lotka-Volterra community model is widely known and provides a phenomenological description of many different communities independent of the specific mechanisms underlying species interactions. Additionally, the Lotka-Volterra model is based on a few simple assumptions, which allows a more general understanding of the phenomenon.

We assumed a two trophic level Lotka-Volterra community model

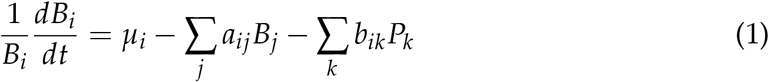

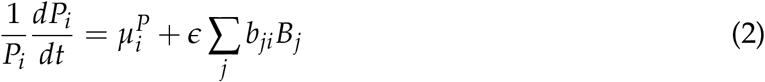

where *B_i_* is the density of prey species *i* with intrinsic growth rate *μ_i_, a_ij_* is the species-specific interaction between prey species *i* and *j*, *b_ik_* is the predation of predator *k* on prey species *i, P_k_* is the density of predator *k* and 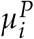 is the mortality rate of the predator. We assumed that there are no direct interactions between predators. *ϵ* is the trophic conversion efficiency between consumption of prey biomass and production of predator biomass; we assumed a trophic efficiency of *ϵ* = 0.1.

We defined the community parameters *μ_i_* and *a_ij_* according to Macarthur & Levins (1967) and Barabás & Meszéna (2009), which specifies a Lotka-Volterra model based on underlying competition of prey species for a resource continuum. Each prey species was identified by a single trait *x_i_*, e.g. body mass, which defined its resource consumption spectrum *u_i_*, i.e. 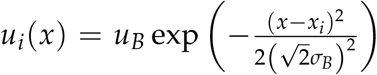, where *x* is the resource identity, e.g. body mass of the resource, *σ_B_* is the niche breath and *u_B_* is a normalizing constant. The competitive interaction between two prey species *i* and *j* is given by 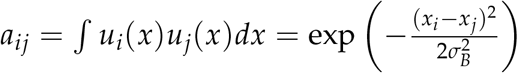, and the scaling constant *u_B_* was chosen such that *a_ii_* = 1 (Barabás & Meszéna, 2009). The intrinsic growth rate *μ_i_* depended on the carrying capacity of the resource *R*(*x*), which we assumed to be a Gaussian, i.e., 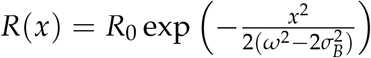, where *R*_0_ is the maximum resource availability and *ω* is the breath of the resource axis, such that 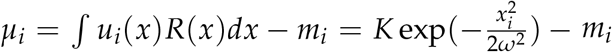, where *K* denotes the maximum intrinsic growth rate, *m_i_* = 0.1 is the mortality rate and *ω* is the niche breadth.

We also assumed a Gaussian predation kernel for the predators. Each predator species was defined by a single trait *y_j_* for predator species *j* which defined its predation preferences. Predation coefficients were given by 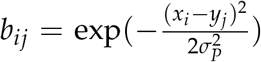 is the niche breath of the predator. Finally, we assumed that all predators have the same mortality rate 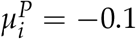.

Community assembly consisted of four steps:

1. Generate a random invader: This invader has a random trait location 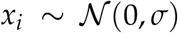 and is either a prey or a predator species. 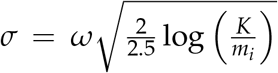 was chosen such that about 98% of the invading prey species had a positive intrinsic growth rate.
2. Compute the invasion growth rate of the invader: The invasion growth rate is defined as 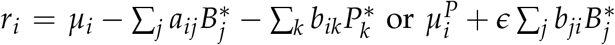, depending on the trophic level of the invader, where 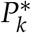 and 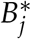 are the equilibrium densities of the current community. If *r_i_* is negative the invasion fails and we move to the next time step.
3. Test feasibility: Compute the new equilibrium of the invader plus the resident species, and if all species have positive equilibrium densities the invader successfully invaded and we move to the next time step. If one species has negative equilibrium density move to step 4.
4. Find new resident community: If the invader replaced at least one species we need to find the new resident community. We computed the equilibrium of all possible sub-communities and removed all non-feasible sub-communities. For the feasible sub-communities we computed the invasion growth rates of the non-present species. If all non-present species have a negative invasion growth rate, the community is saturated. To determine the next resident community, we selected the most species-rich, feasible, saturated sub-community. If there were multiple communities of equivalent richness, we randomly selected one. This method of determining the new resident community led to the same qualitative dynamics as introducing each invader at low densities in the model, and simulating the community dynamics until equilibrium was reached (Appendix S3, Figure S3)

In the main text we focus on a simplified version of community assembly which assumes that the time between invasions was sufficiently large that the community would reach an ecological equilibrium between invasions (Serván & Allesina, 2021). Additionally, we ignored transient dynamics as well as potential non-equilibrium behavior (Serván & Allesina, 2021; Law & Morton, 1996). In the Appendix we show that these simplifications do not affect our main conclusions (Appendix S3).

## Results

We simulated community assembly under two different conditions, with and without predators present (Fig. 1). Without predators, there was exactly one stable configuration of prey species, and the trait distance between prey species was roughly twice the niche breath of the prey species, i.e. 2*σ_B_* (Macarthur & Levins, 1967; Barabás *et al*., 2012). Community assembly always converged towards this single stable configuration, independent of the invasion history, which aligns with previous theoretical predictions (MacArthur, 1970). Over time, the probability of successful invasion by a new arrival decreased towards zero (blue shaded area, Fig. 1 A). Overall, results for the one trophic level community model are consistent with the expectation of convergence towards a stable endpoint known from previous models.

**Figure 1:**
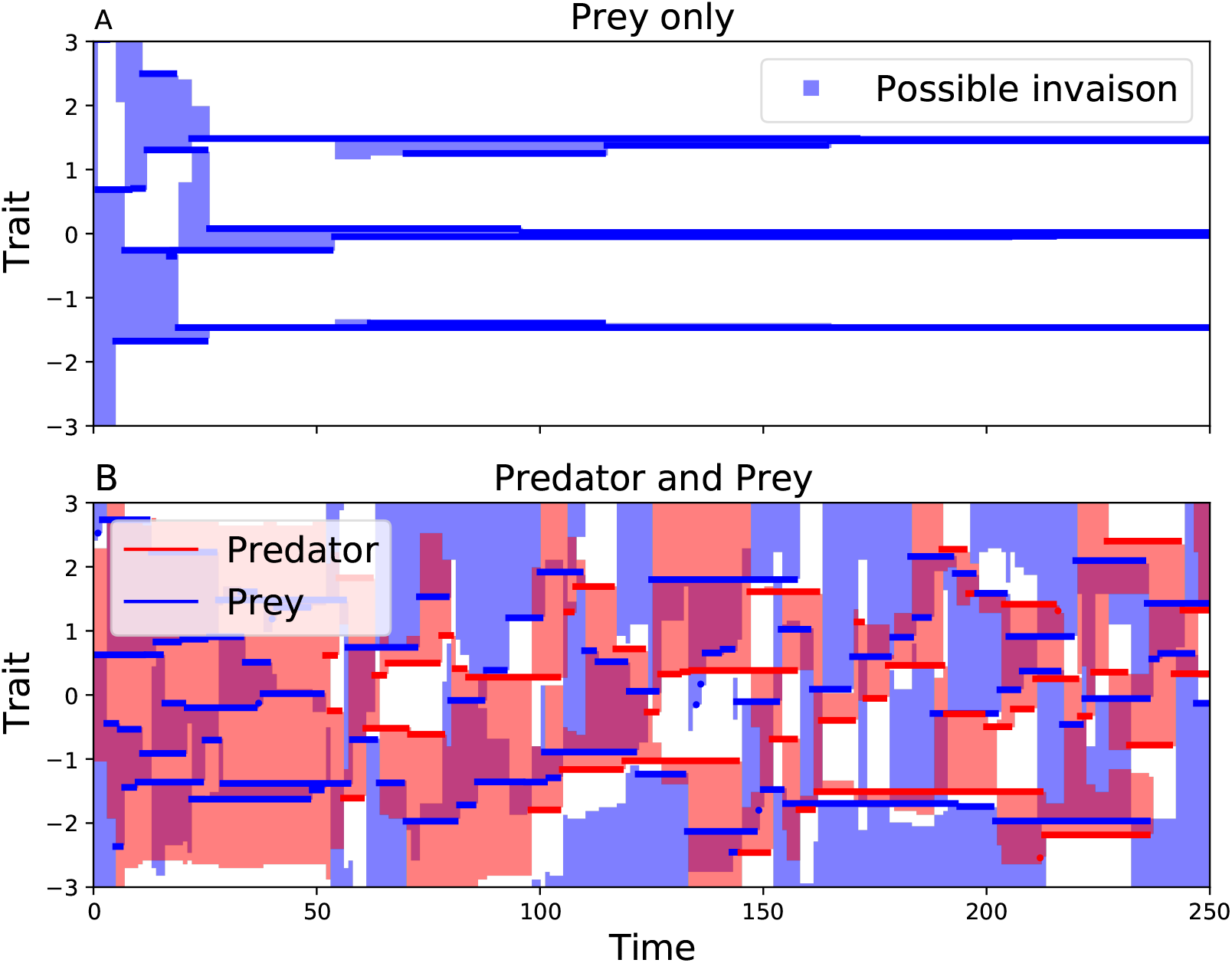
We simulated community assembly in one- (A) and two-trophic-level (B) communities. Each year (x-axis) a new species with a random trait (y-axis) is introduced to the community and potentially replaces residents. A: In the absence of predators, the prey species self-organize into a regular pattern known as limiting similarity. This final composition is stable and does not depend on the community assembly process. B: The inclusion of predators changes the community assembly from being deterministic and stable to unpredictable, characterized by continuous invasions and extinctions. There is no stable, uninvadable configuration. Shaded regions indicate trait values for which a potential invader would be successful. Without predators, these regions disappear over time. Conversely, in the presence of predators, invasion by a prey species tends to increase the potential for invasion by predator species, and vice versa.

The inclusion of a second trophic level qualitatively changed the dynamics. The two trophic level model did not lead to the typical trait distribution known from limiting similarity, with equally spaced species at a few unchanging trait values (fig. 1A). Rather, the two trophic level community exhibited continuous invasion of new species and extinction of established species, although with no trend in species richness. A late-arriving species did not have a lower probability of invasion success than an early-arriving one. Consequently, community assembly was not directed, and did not converge towards a stable end point.

Intuitively, we can understand this continuous invasion and extinction by considering an example with just two prey species, B_1_ and B_2_, and two predator species, P_1_ and P_2_ (Schreiber & Rittenhouse, 2004). We assume that P_1_ is a better predator for B_1_ and P_2_ a better predator for species B_2_; the predators are equivalent in all other aspects. Given the community composition (B_1_, P_1_), the prey species B_1_ has low fitness because of strong predation pressure from P_1_. Therefore, prey species B_2_ can invade and exclude B_1_, leading to the community (B_2_,P_1_). However, P_2_ is a better predator for B_2_ and will consequently displace P_1_, leading to the community (B_2_, P_2_). Under these conditions B_2_ will have low fitness because of strong predation pressure from P_2_, therefore B_1_ will invade leading to (B_1_,P_2_). Finally, to close the cycle, P_1_ will replace P_2_ as it is a superior predator for species B_1_. Our model was driven by qualitatively similar dynamics, though the randomness in the traits of potential invaders complicates the simple cycle.

This cycle depends on sufficiently specialized predators such that the community (B_1_, B_2_, P_1_) is not stable (Schreiber & Rittenhouse, 2004). In our simulations, this meant that the niche width of the predator *σ_P_* had to be smaller than the niche width of the prey species *σ*_B_ (Fig. 2). Results from limiting similarity theory give us an intuitive understanding of this condition. From limiting similarity we expect the coexisting species to be separated by roughly twice their niche breath, i.e. 2*σ* (Macarthur & Levins, 1967). This result is quite robust to changes in the fitness function and the competition kernel (Barabás *et al*., 2012). Let Ω denote the length of the interval of feasible trait values for prey species, i.e. 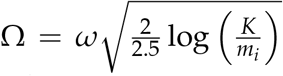, which is also roughly the interval of feasible trait values for predator species. Then we expect ~ Ω/2*σ_B_* prey species and ~ Ω/2*σ_P_* predator species in a stable configuration. However, at stable equilibrium the number of predator species cannot exceed the number of prey species (Tilman *et al*., 1982; Meszéna *et al*., 2006). We therefore conclude that a stable configuration implies *σ_P_* ≥ *σ_B_* (see Appendix S2 for a more precise proof). Note however, this argument only tells us that we should not expect a stable configuration for *σ_P_* < *σ_B_*, it does not necessarily imply that we should expect a stable configurations for *σ_P_* > *σ_B_*.

**Figure 2:**
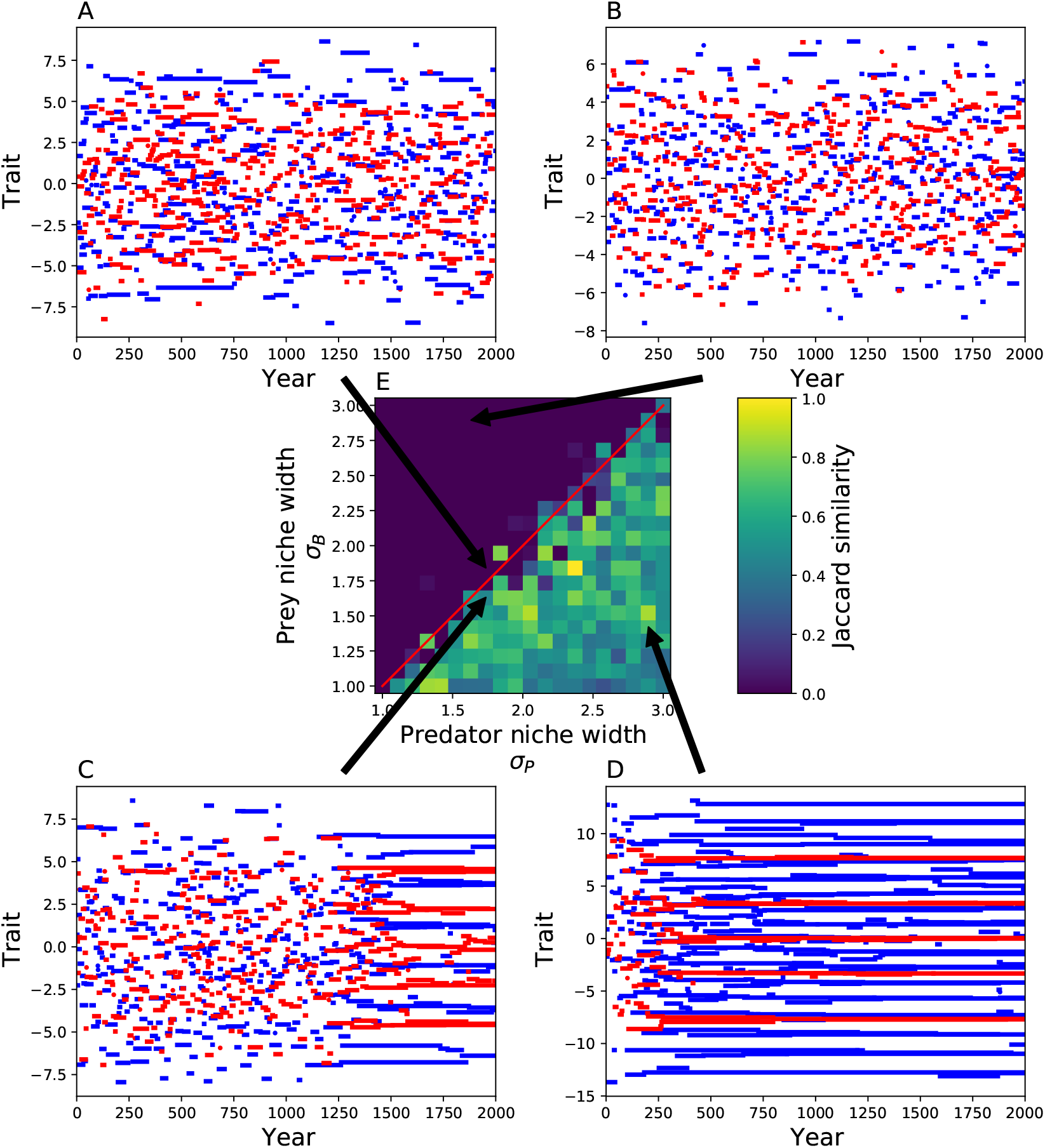
We simulated the two-trophic community with different values of niche width for predator and prey communities for 2000 invasion cycles. A-D show examples of community dynamics, the arrows show to the corresponding niche width values. E: We report the Jaccard similarity of the community at the end and the community 200 steps before the end point. Communities with higher prey niche width (y-axis) than predator niche width (x-axis) show continuous assembly patterns (e.g. Panel A and B). Conversely, communities with higher predator niche width converge towards a stable community (Panel C and D). The environmental niche breath was chosen as *ω* = 3*σ_P_* to avoid boundary issues (Appendix S2).

### Stability within the disorder

The two trophic level community model led to unpredictable assembly, meaning that community composition cannot be predicted far into the future. In contrast, the trait distribution of the community (the number of species with traits in a particular interval of trait values) remained largely unchanged (Fig. 3). Typically, an invader replaced a resident species with a similar trait, as the invader’s presence has the largest effect on similar species (Vannette & Fukami (2014) and Fig. 3 A, D). Consequently, each individual invasion had no large effect on the trait distribution. On a longer time scale, the prey species used essentially all available resources: if a certain range of the resource spectrum remained unused, then an invader soon filled this gap. As a result, the trait distribution of the prey species mimicked the underlying resource distribution (MacArthur, 1970), which was constant over time.

**Figure 3:**
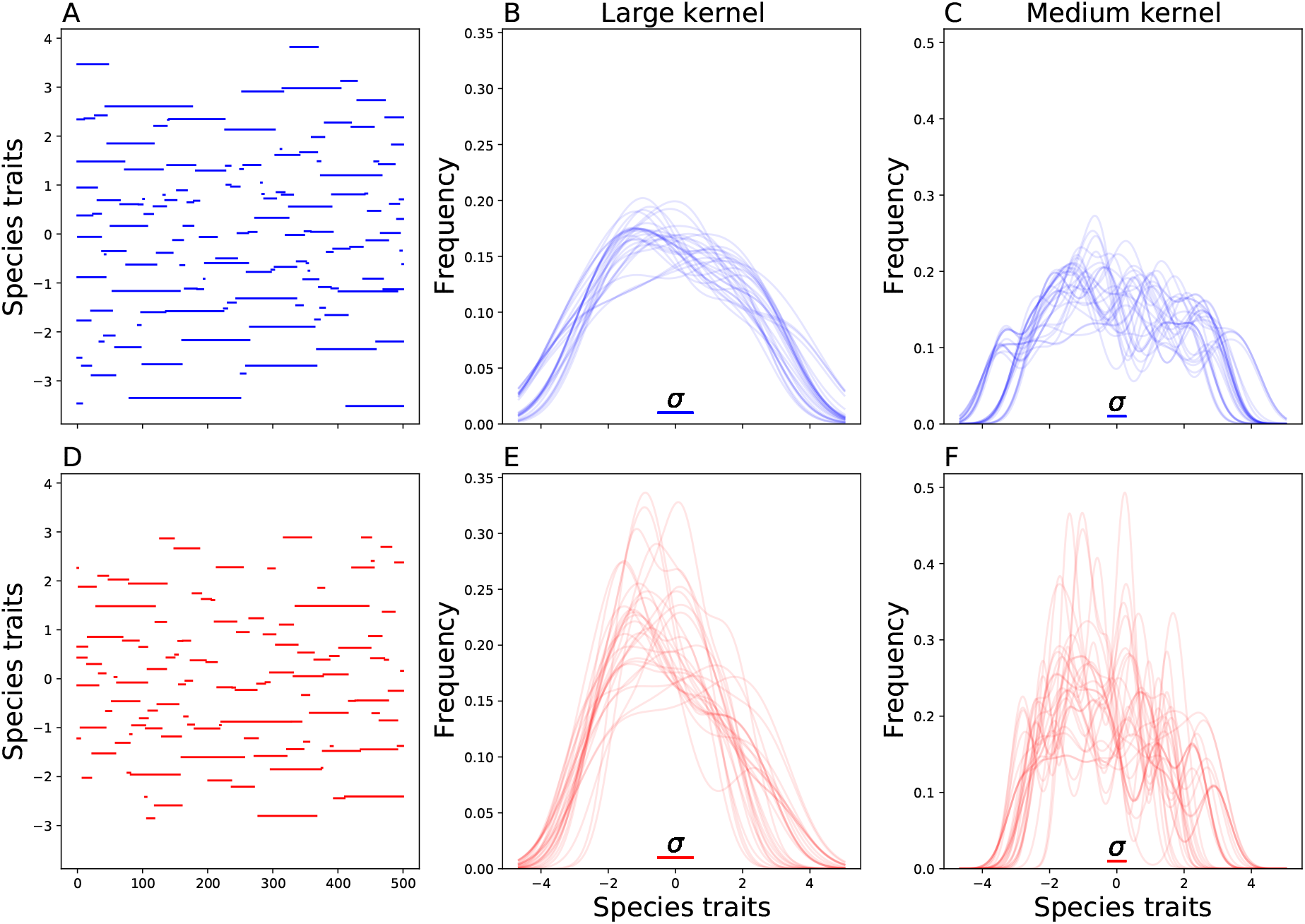
A,D: The specific species composition at each time-point is stochastic and changes very fast. B,E: We computed the trait distribution with a Gaussian kernel density estimate, the kernel size is shown with the inlet. Each line corresponds to a given time point. The resulting trait distribution is largely stable for both predator and prey species. D,F: The resulting trait distribution is less stable at a smaller kernel size. Generally, we expect the trait distribution to be roughly stable if the kernel size corresponds to the competition kernel of the species.

Similarly, the trait distribution of the predator species was roughly constant, albeit more variable over time than the trait distributions of prey species. Intuitively, the predator trait distribution mimicked their underlying resource distribution, i.e. the abundance of prey species. However, this underlying resource distribution was not perfectly constant, but rather varied slightly over time. The trait distribution of the predator species is therefore a roughly constant approximation of the underlying roughly constant trait distribution of the prey species.

A consequence of this stable trait distribution was the over-dispersion of species traits compared to a randomly selected community without competitive interactions (Fig. 4, C and D). Although we did not observe any strict lower limit to the trait difference between two coexisting competing species, we rarely observed coexisting species with very similar traits.

**Figure 4:**
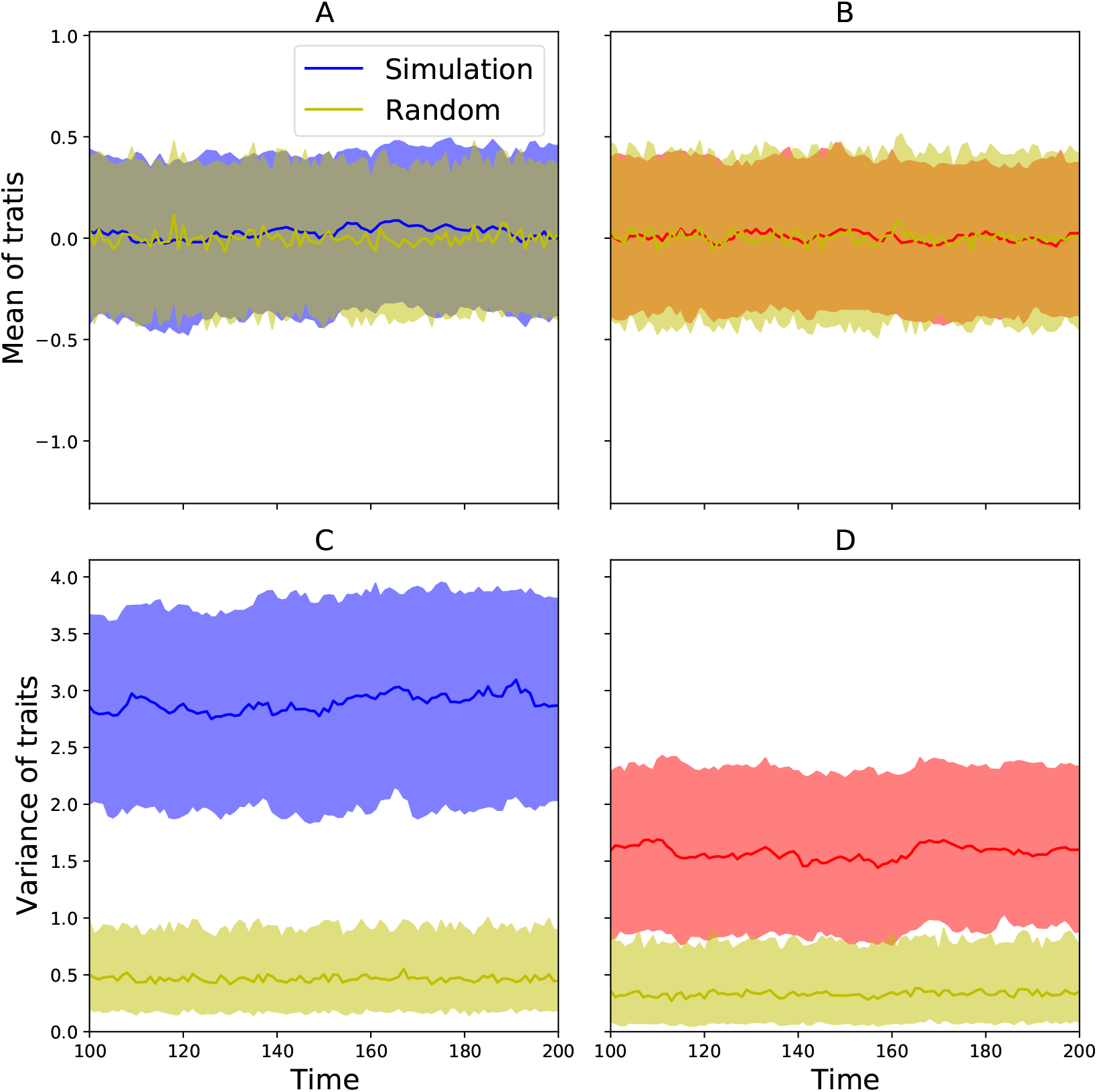
We compared the trait distribution resulting from the community assembly (blue for prey species [left column] and red for predator species [right column]) to distributions from random species selection (yellow). A,B: The trait mean from the community assembly did not differ from a randomly selected sample of species. C,D: However, the species traits were overdispersed over the available trait space, compared to randomly selected species. A-D: Lines show median across replicate simulations, shaded areas show 25-75% percentile lines.

## Discussion

Our paper highlights the idea that community assembly does not always move towards a stable endpoint, but rather that communities can remain indefinitely in transient-like behavior with high species turnover. For such communities, the term community assembly is somewhat inappropriate, as there is no final community to be assembled. Our modeling results make it clear that this “continuous assembly” dynamic depends only on the presence of sufficiently specialized predators (Fig. 1, Appendix S2). The open question then is how widespread we should expect such dynamics to be in nature.

Whether this mechanism is actually present in natural communities is currently difficult to answer, as three conceptually different mechanisms can lead to the continuous assembly observed in nature (Dornelas *et al*., 2019). Specifically, continuous assembly can be driven by external environmental changes (Dornelas *et al*., 2019), stochastic fluctuations based on neutral dynamics (Hubbell, 2001) or internal dynamics as described here. Yet, these different underlying mechanisms lead to different links between invasion and extinction events. In the neutral model invasion and extinctions are independent of each other. In the case of external environmental change, the invasion and extinction are not causally linked but are both driven by the same external factor. We would therefore expect a correlation, but no causal link. Finally, in the case of internal dynamics, invasions cause extinctions and vice-versa, and we would therefore expect a causal link as well as a positive correlation.

The BioTIME data set offers a possibility to assess whether invasions and extinctions are correlated and potentially linked. As a cursory analysis, we investigated the correlations between invasions and extinctions in the BioTIME data (Appendix S1). We found that in 24 of the 44 datasets (~ 55%), the observed correlation was significantly higher than expected by chance, i.e., *p* < 0.05 (Fig 5). For 17 of the 44 datasets (~ 40%), the observed correlation was stronger than any correlations found in 1000 randomizations. Aquatic ecosystems in particular showed a stronger correlation than expected by chance. Interestingly, Li *et al*. (2022) found that predation kernel width scales differently with body size in aquatic ecosystems compared to terrestrial ecosystems, which potentially explains why continuous assembly is more frequent in aquatic communities. The strong correlation between invasion and extinctions is consistent with either internal or externally driven invasion and extinction, however we were not able to test whether there is indeed a causal link between invasions and extinctions. Additionally, one might test whether invasion and extinction events correlate with strong changes in environmental factors to understand whether and which external factors drive this continuous community assembly.

**Figure 5:**
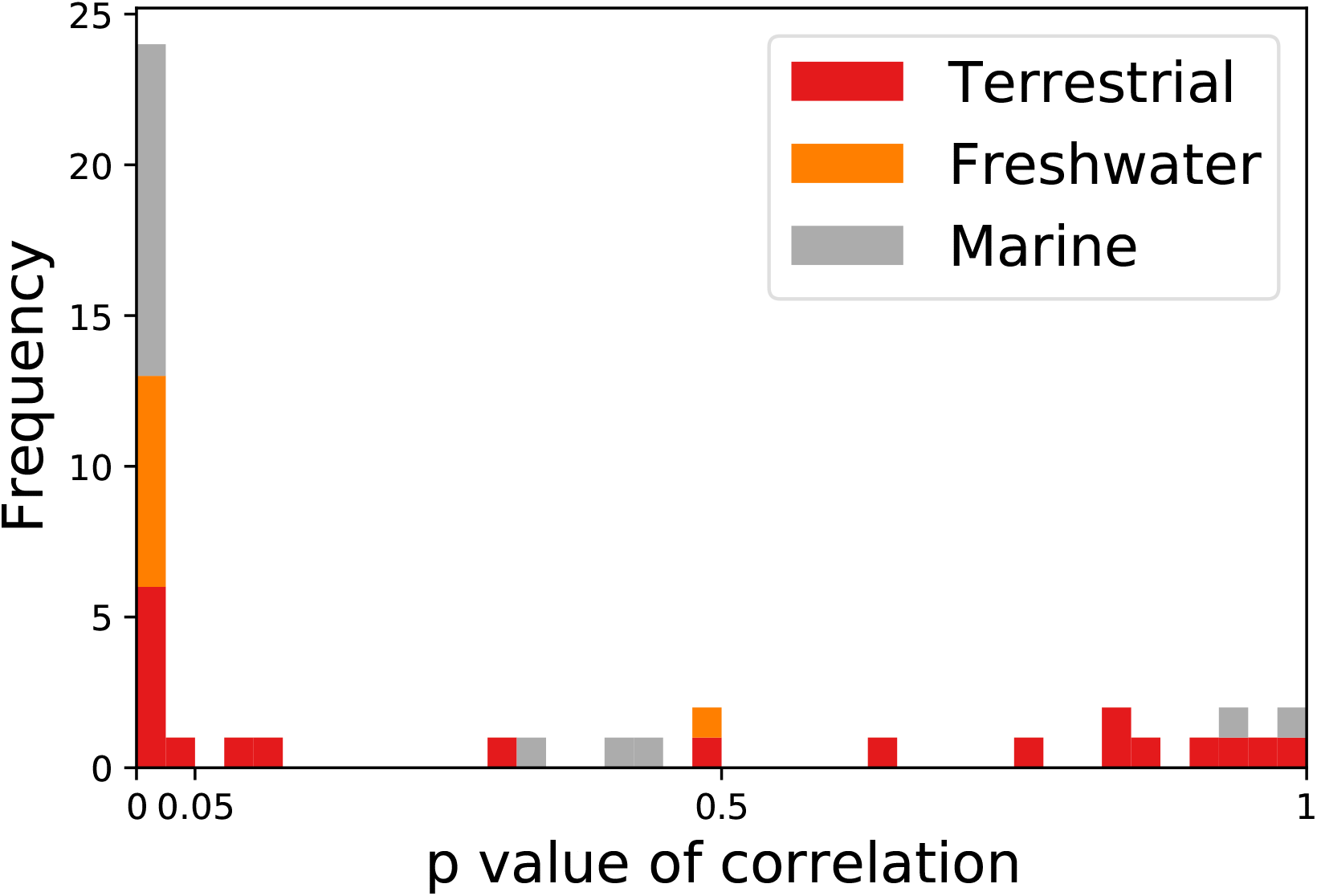
We report the p-value of the correlation between invasions and extinctions observed in the BioTIME datasets compared to correlations based on random rearrangements of the years in each dataset. For around 50% of the data sets the correlation was significantly higher than expected by chance, as expected from theory. This pattern appears to be driven by freshwater and marine communities.

### Why do we see continuous assembly?

Our first research question focused on the conditions necessary for continuous assembly to emerge in our model. We found that sufficient specialization of the predators was the key condition (Schreiber & Rittenhouse, 2004, Appendix S2), because it allows prey species to competitively exclude other prey while not sharing their predators. To understand this dynamic intuitively, we observe that a prey species with no specialist predator will have high fitness, allowing it to reach high abundance and displace competitors with similar traits. However, as the prey species reaches high abundance, a niche is created for a predator with the corresponding trait to invade. The predator then reduces the prey species’ fitness and abundance, opening the possibility for other prey species with similar traits to invade. If predators are sufficiently specialized, some of these new invading prey species will not experience high predation pressure and will have high fitness. Predators do not drive the prey species to extinction directly, rather they reduce the fitness of their prey to the point where they can no longer compete with neighboring prey species that experience far less predation pressure.

Abrams & Matsuda (1997) described a similar pattern of continuous assembly in evolutionary dynamics. They investigated a community with two prey species B_1_ and B_2_ and one predator which alters its predation preference either through evolution or behavioral changes. Whenever a prey species becomes abundant the predator shifts its preference towards this prey species, reducing its abundance. The other prey species, without any predation pressure, becomes abundant until the predator switches its preference again. Essentially, the predator is chasing the food in the trait-space. In our model, the same dynamics drive continuous assembly, though individual predators do not change their predation preferences, but rather a new predator invades the community.

We emphasize that building a model capable of producing continuous assembly is relatively easy. Continuous assembly has emerged independently in several different community models of various complexity, including our two-trophic Lotka-Volterra model, a size based predation model (Law & Morton, 1993; Morton & Law, 1997), a two-trophic level mechanistic resource competition model based on empirical plankton traits (Spaak *et al*., 2022), various food-web models (Hamm & Drossel, 2021; Allhoff *et al*., 2015; Loeuille & Loreau, 2005) and Lotka-Volterra community models with random species interactions (Bunin, 2017; Barbier *et al*., 2018). In addition, the evolutionary dynamics of Abrams & Matsuda (1997) have been confirmed in other theoretical models (Cortez & Ellner, 2010; Cortez, 2016) and empirical observations (Becks *et al*., 2010). To our knowledge, none of these investigators designed their model to produce continuous assembly – it emerged on its own, indicating that the phenomenon is robust to different model assumptions.

Continuous assembly has also been described in Lotka-Volterra community models with random species interaction matrices (Bunin, 2017), but the underlying mechanism appears to be different. The random interaction models show continuous community assembly if the circle containing the eigenvalues of the interaction coefficients matrix approaches 0, which is related to the variance of inter-specific interaction strengths (Bunin, 2017) and conceptually similar to the classical diversity-stability debate (May, 1972; Allesina & Tang, 2012). However, the eigenvalue distributions for the two trophic level community model investigated here are not similar to the eigenvalue distributions of random interaction coefficient matrices.

### What remains stable in this disorder?

While the continuous assembly process leads to unpredictability in community composition in our model, we found that species richness and the trait distribution remained largely constant over time (Fig. 3, Appendix S4, Figure S5). The stable trait distribution matches experimental findings from Goldford *et al*. (2018), who assembled multiple microbial communities and found large differences in species composition among replicates. However, the relative abundance of taxonomic families remained largely constant across different replicates, similar to how the trait distribution remained constant in our model. Similarly, the fraction of predator species remained relatively stable, despite the continuous turnover of species. This matches findings from food-web models which found a continuous community assembly, but relatively stable trophic level distributions (Hamm & Drossel, 2021; Allhoff *et al*., 2015).

### Limitations and future work

Our theoretical model predicts that the relative niche breadth of the trophic levels have strong implications for the stability of the emerging community. But what does this mean in a natural community? In building our model we imagined a trophic food-web where predation is driven by body-size, e.g. zooplankton as the prey species and small fish as the predator species (Hamm & Drossel, 2021; Allhoff *et al*., 2015; Williams & Martinez, 2000). In this context, different niche breadth implies that the fish consume a smaller range of different zooplankton body-sizes than the range of different phytoplankton body-sizes the zooplankton consume. Unfortunately, we do not know whether higher trophic levels actually are more specialized than lower trophic levels. Li et al. (2022) analyzed the ratio of predator to prey body masses and found that larger species tend to have slightly wider niches than smaller species. However, Li et al. (2022) analyzed link probability and did not include any information about link strength. Additionally, they focused on the effect of predator body size on niche width, and not how trophic status itself affects niche width, although trophic status and body size are generally well correlated (Riede *et al*., 2011). What niche breadth implies in a context of plants and herbivores is less clear. Perhaps it means that herbivore diets have tighter stochiometric constraints than plant resource requirements. On the other hand, we know of many specialist predators and pathogens (Bever *et al*., 2012) which might promote a continuous assembly pattern (Schreiber & Rittenhouse, 2004).

Our model was relatively simple, allowing only for two-trophic levels and no omnivory or cannibalism, which is widespread in natural communities (Williams & Martinez, 2000; Allhoff *et al*., 2015). It would be interesting to see whether our findings apply to more complex niche-based food-webs. Currently food-web models typically assume that predation kernels are independent of trophic status or body size (Loeuille & Loreau, 2005; Emmerson & Raffaelli, 2004; Allhoff *et al*., 2015; Hamm & Drossel, 2021; Williams & Martinez, 2000; Brose, 2010). That is, these models assume *σ_B_* = *σ_P_*, which is exactly what we have identified as the boundary between continuous assembly and stable equilibria. This potentially explains why some of these show a pattern of continuous assembly (Hamm & Drossel, 2021; Allhoff *et al*., 2015), while others show stable community compositions (Loeuille & Loreau, 2005). However, the models also differ in other aspects, such as the response function to predation or the number of traits per species. It is currently unclear which of these model differences affect the community assembly process.

If the changes in community composition observed in natural communities are indeed driven by internal mechanisms as described here, then we would have to reconsider core concepts of community ecology which are based on equilibrium dynamics. Specifically, modern coexistence theory and its dependence on invasion growth rates into stable equilibrium dynamics (Ellner *et al*., 2019; Spaak & De Laender, 2020; Barabás *et al*., 2018), ecosystem stability based on linearization of the community dynamics near the equilibrium (May, 1972; Allesina & Tang, 2012, 2015) and potentially biodiversity ecosystem-function relationships, which are typically evaluated after the community has fully assembled (Bannar-Martin *et al*., 2018; De Laender *et al*., 2016).

## Acknowledgments

We thank Matthieu Barbier for his insightful comments and our understanding of continuous invasion in random matrices. J.W.S was supported by the Swiss national science foundation SNSF under the project P2SKP3 194960. S.P.E. was supported by US NSF grant DEB-1933497. P.B.A. was supported by US NSF grant DEB-1933561.

## Appendix

### S1 BioTIME data

We compared the patterns from the simulations to patterns in empirical data from BioTIME, a database of community assemblage time-series across the world (Dornelas *et al*., 2018). We focused on presence-absence patterns and therefore aggregated each time-series to annual scale, i.e. a species was assumed to be present if it was observed at least once in a given year, otherwise it was assumed to be absent. To observe patterns over time, we focused on datasets with at least 30 years of sampling. We found a total of 44 suitable datasets representing different taxonomic groups (birds, fish, invertebrates, terrestrial plants, benthos, mammals and amphibians), different biomes (lakes, rivers, different marine waters, different types of forests and prairies) with latitude ranging from 62.1° south to 67.1° north. The species richness ranged from 1 to 2000 per year and from 6 to 4120 over the respective observation periods.

We computed species richness, the proportion of invasions, and the proportion of extinctions per year for each dataset. The proportion of invaders in year *t* was defined as the number of species present in year *t* which were not present in year *t* – 1 divided by the species richness in year *t*. Similarly, the proportion of extinctions in year *t* was the number of species present in year *t* but not in year *t* + 1 divided by the species richness in year *t*. We then computed the correlation between the proportion of invaders in year *t* and the proportion of species going extinct between year *t* and *t* – 1. We compared the observed correlation of each dataset to the correlation of invasions and extinctions in the same dataset if the years were randomly reshuffled. We report the p-value of observing a correlation as high or higher than 1000 randomizations.

### S2 Stable communities

In the main text we have focused on the cases where the two trophic levels lead to continuous changes in community composition. Generally, this is observed to be the case if the niche width of the predator is smaller than the niche width of the prey. If the niche width of the predator is sufficiently large then a stable community is possible (Fig. 2). Note that “stable community” here means both internal and external stability: the species that are present are coexisting at a locally stable equilibrium, and no potential invader has a positive invasion growth rate.

In this section we prove that a stable community cannot occur if *σ_P_* is sufficiently small compared to *σ_B_*, under two additional assumptions.

1. The prey species are evenly spaced at some distance *D_B_*.
2. The niche space is very large, i.e. *ω* ≫ *σ_B_*, and effectively infinite (the precise meaning of “effectively infinite” will be clarified below).
3. The consumption kernels of the predators are sufficiently narrow that each predator effectively consumes only one prey species, i.e. *σ_P_* is small relative to *D_B_*.

In our simulations, stable communities (when they occur) always have prey species evenly spaced, except near the boundary of the niche space. Other theoretical studies (Szabó & Meszéna, 2006; Macarthur & Levins, 1967; Barabás *et al*., 2012) have also generally found that species are evenly spaced, for a wide range of intrinsic growth rates and competition kernels. When the trait space is unbounded, rescaling the niche axis relative to *σ_B_* implies that the equilibrium prey spacing *D_B_* is proportional to *σ_B_*. Assuming that *σ_P_* is small relative to *D_B_* is thus equivalent to assuming that *σ_P_* is small compared to *σ_B_*. In our simulations, stability ceases to occur when *σ_P_* is only slightly smaller than *σ*_B_, but our arguments here only show that stability is impossible when *σ_P_* is considerably smaller than *σ*_B_.

Assumption 2 implies that all prey species have identical intrinsic growth rates. Mathematically, we will use assumption 3 to show that each predator species must be located “on top of” a prey species (i.e., it must have the same trait value as one of the prey species). Assumption 1 will be used to show that a prey with a predator directly on top of it can be invaded by a prey species with a slightly similar trait value, hence a community with that feature cannot be stable. These two properties together imply that a stable community cannot occur.

Importantly, without assumption 2, stable communities are possible even when *σ_P_* is small. Specifically, if we assume *ω* < *σ_P_* < *σ_B_*, then one example of a stable consists of exactly one predator and one prey species, both having trait value 0 (Fig. S1).

#### S2.1 Predators cannot be located between prey species in a stable community

We show that each predator in a stable configuration must have a prey with identical trait. We prove this by assuming that a predator *j* exists with trait *x_j_*, and the closest prey species has trait *x*_0_ = *x_j_*. Then the growth rate of the predator, which must by assumption be 0, is

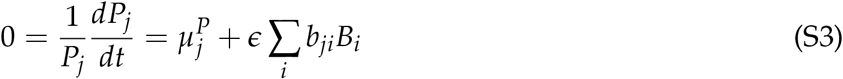

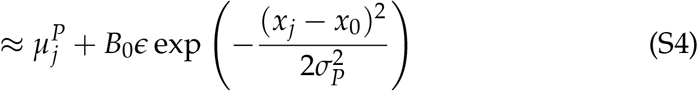

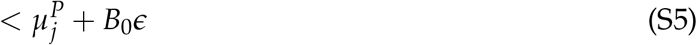

**Figure S1:**
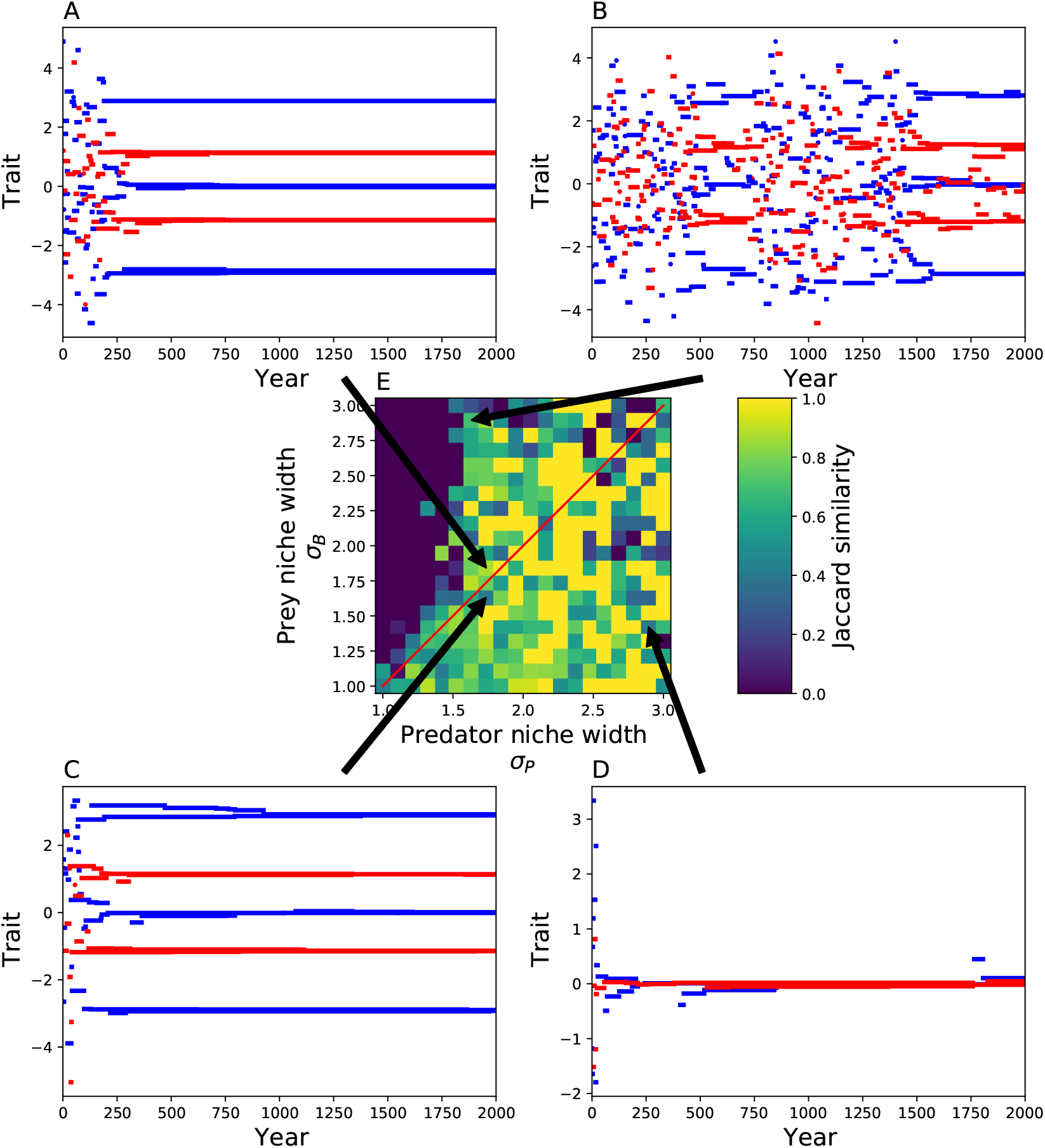
Similar to figure 2 we simulated the two trophic level community with different values of niche width for predator and prey communities for 2000 invasion cycles. However, we here chose a fixed environmental niche width of *ω* = 2.5. Because the predator niche width *σ_P_* is comparable to the environmental niche width *ω* communities can be stable despite *σ_P_* < *σ_B_*, e.g. panel B. A-D show examples of community dynamics, the arrows show to the corresponding niche width values. E: We report the Jaccard similarity of the community at the end and the community 200 steps before the end point.

From S3 to S4 we used the fact that prey species are equally spaced at distance *D_B_*, and the consumption rates of the predator *j* on all other prey species are therefore 0. Equation S5 then shows that a invading predator with trait *x* = *x*_0_ would have a positive growth rate, so the the system is therefore not stable. Consequentially, each predator in a stable community is located exactly on top of a prey species.

From the same calculation it follows that if a predator has the identical trait as a prey species, there can be no other predator *j′* with trait value consuming the same prey species, for any such predator would have a negative population growth rate.

This leads to two additional insights for situations where *ω* is large compared to *σ_B_* but finite, and *σ_P_* is small compared to *σ_B_*:

1. Each prey close to the center of the niche space has a predator with identical trait and all prey species close to the center of the niche have identical equilibrium abundance.
2. All predators at the center of the niche have identical equilibrium abundance and are also equally spaced with distance *D_B_*.

Note, this does not correspond to the stable systems observed in figure 2 because in those communities we do not have a small *σ_P_*. For large *σ_P_* a predator can (and will) have a trait value between prey species.

#### S2.2 Predators cannot be located on prey species in a stable community

As shown above, in any stable community with sufficiently small *σ_P_* we must have equidistantly spaced prey and predator species, separated by distance *D_B_*. Without loss of generality we can assume that one of the prey species has trait value *x* = 0 (i.e., we pick one prey species, and measure traits relative to that of the chosen species). We will show that a species with some trait *x* = *ε* with |*ε*| ≪ 1 has a positive invasion growth rate, therefore the community is actually not stable. This shows that a stable community cannot actually exist.

Let *r*(*ε*) denote the invasion growth rate of a species with trait *ε* very close to 0. We must have *r*(0) = 0, as the species with trait value 0 is at equilibrium. Further, if *r′*(0) ≠ 0 at *x* = 0, then *r*(*x*) is positive for some *x* ≈ 0, implying a nonstable community. So it suffices to show that the second derivative *r″*(0) is positive, because when *r′*(0) = 0 the second-order Taylor series 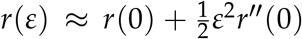 implies that *r*(*ε*) > 0 for sufficiently small *ε* when *r″*(0) > 0.

Prey invasion growth rate in general is *K* – ∑*_j_a_ij_B_j_* – ∑*_k_ b_ik_ P_k_*, so the second derivative is

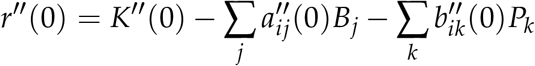

where •″ indicates the second derivative of the interaction coefficients with respect to trait *x*, evaluated at *x* = 0. Note that *K*″ → 0 as *ω* → ∞ because *K* becomes constant; here we specify that *ω* is “effectively infinite” in the sense that *K*″ (0) is small relative to the other terms and can be neglected in calculating *r″* (0).

To evaluate the second derivatives we differentiate the Gaussian kernel twice,

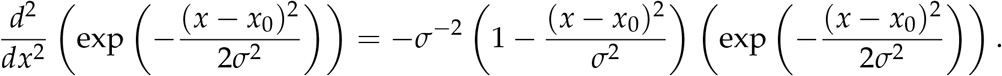

We therefore have (with all sums running over all species in the community)

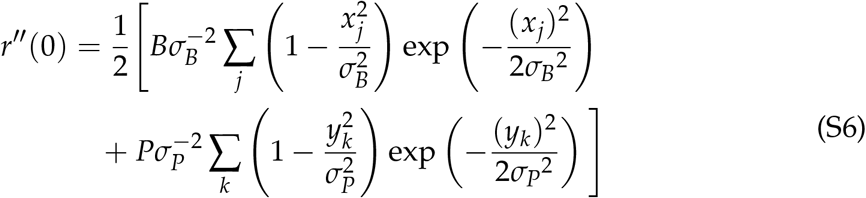

The right-hand side in (S6) is positive when the sums run over the set 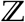 of all integers; this follows the fact that

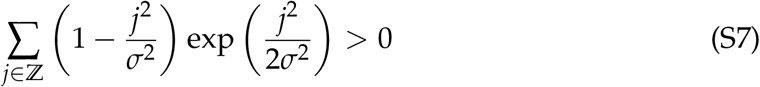

which we will prove below, and the fact that prey and predators occur at trait values ±*jD_B_, j* = 0,1,2,…. We now specify that *ω* is “effectively infinite” in the sense that the set of species in the community (equally-spaced prey and predators, across some symmetric neighborhood around 0) is broad enough that the sign of (S6) is already positive when the sums run over all species in the community, as it is when the sums run of 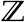. This implies *r″* (0) > 0, so the only possible stable community is in fact not stable.

To finish, we now prove (S7), using the Poisson summation formula

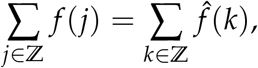

where 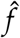 is the Fourier transform of *f*, i.e. 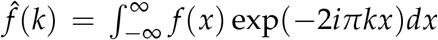, where *i* is not an index but rather 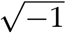. We now compute the Fourier transform of 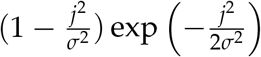 as follows:

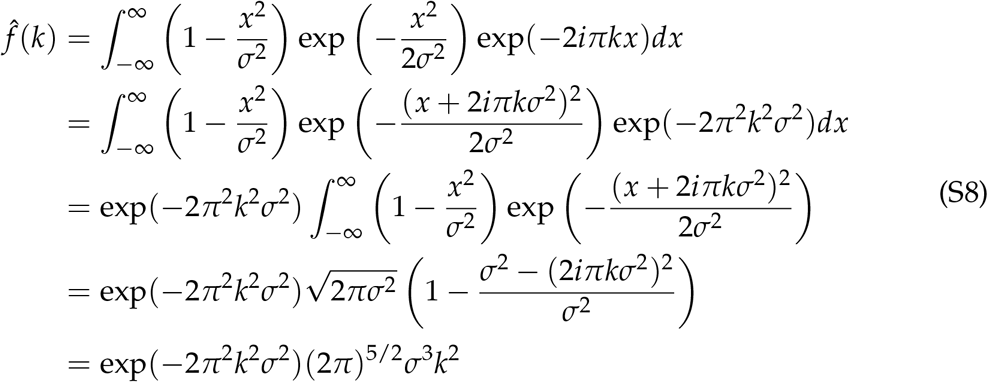

This last expression is positive for all *k*, therefore the sum 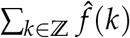 is also positive. The integral was evaluated using the fact that 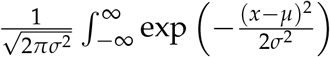 describes a normal distribution with mean *μ* and variance *σ*^2^.

**Table S1:**
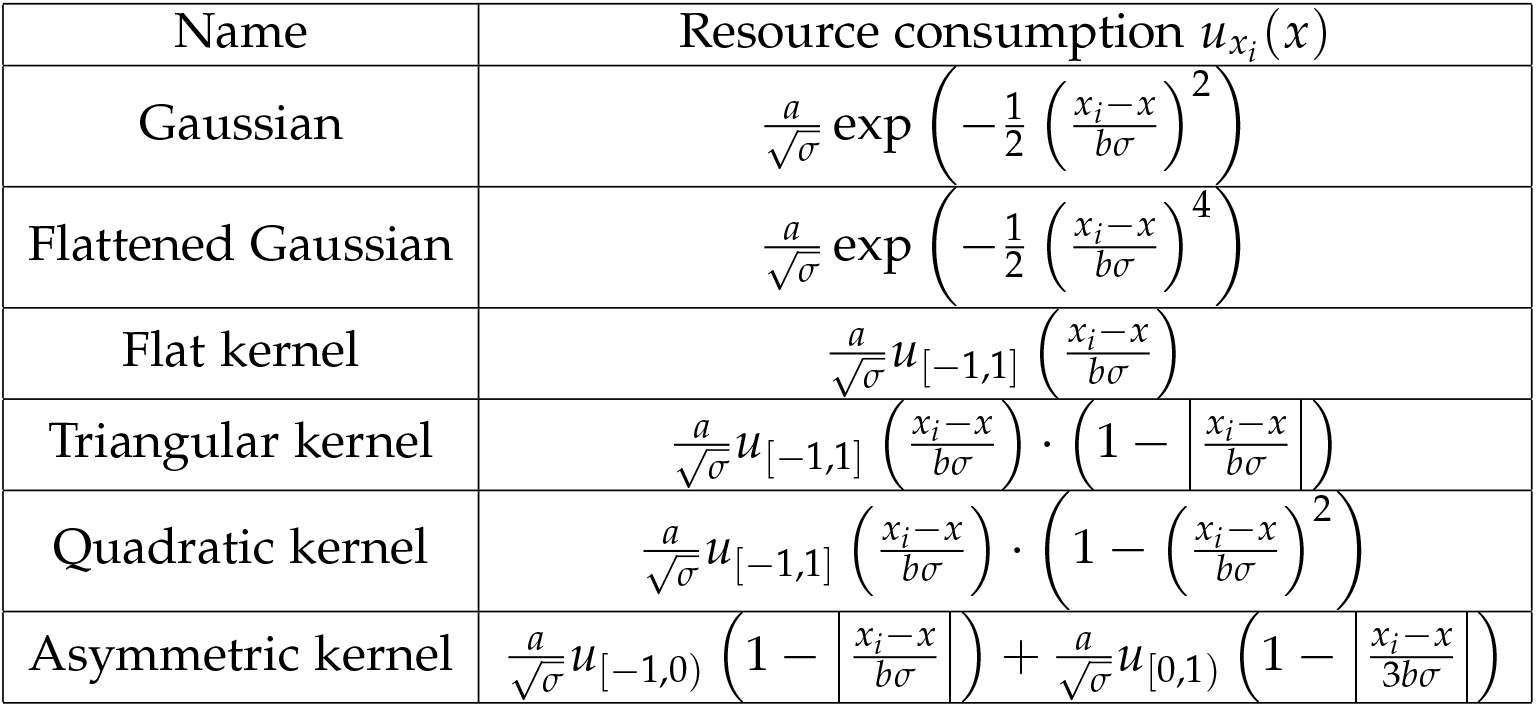
The different competition kernels we have investigated. *u*_[*x*_1_, *y*_2_]_ is the indicator function of the interval [*x*_1_, *x*_2_], i.e. *u*_[*x*_1_, *y*_2_]_(*x*) = 1 if *x*_1_ < *x* < *x*_2_, otherwise it is zero. Fig. S2 shows a visual representation of these kernels. The scaling parameters *a* and *b* are chosen such that 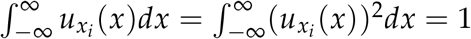.

### S3 More general cases

In the main text we have, for simplicity, focused on specific model. We show here that our main finding, i.e. continuous community assembly, is robust to many different scenarios, including different resource consumption and competition kernels (Fig. S2), simulating population densities over time over time instead of computing the equilibrium dynamics directly (Fig. S3) and a finite regional species pool (Fig. S4).

We investigate a total of six different competition kernels: Gaussian kernel, flattened Gaussian, flat kernel, triangular, quadratic and asymmetric (Table S1). Each resource consumption kernel *u_x_i__*(*x*) is described by the location of maximal resource consumption *x_i_*, the width of the kernel *σ* and two scaling factors *a* and *b*. The coefficient of competition between two prey species with traits *x_i_* and *x_j_* is given by 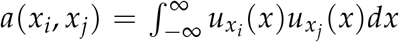. The scaling factors are chosen such that *a*(*x_i_, x_j_*) = 1 and 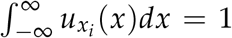, i.e. the kernel only affects the shape of the competition, not however how strong intraspecific competition is, nor how much a predator consumes in total.

**Figure S2:**
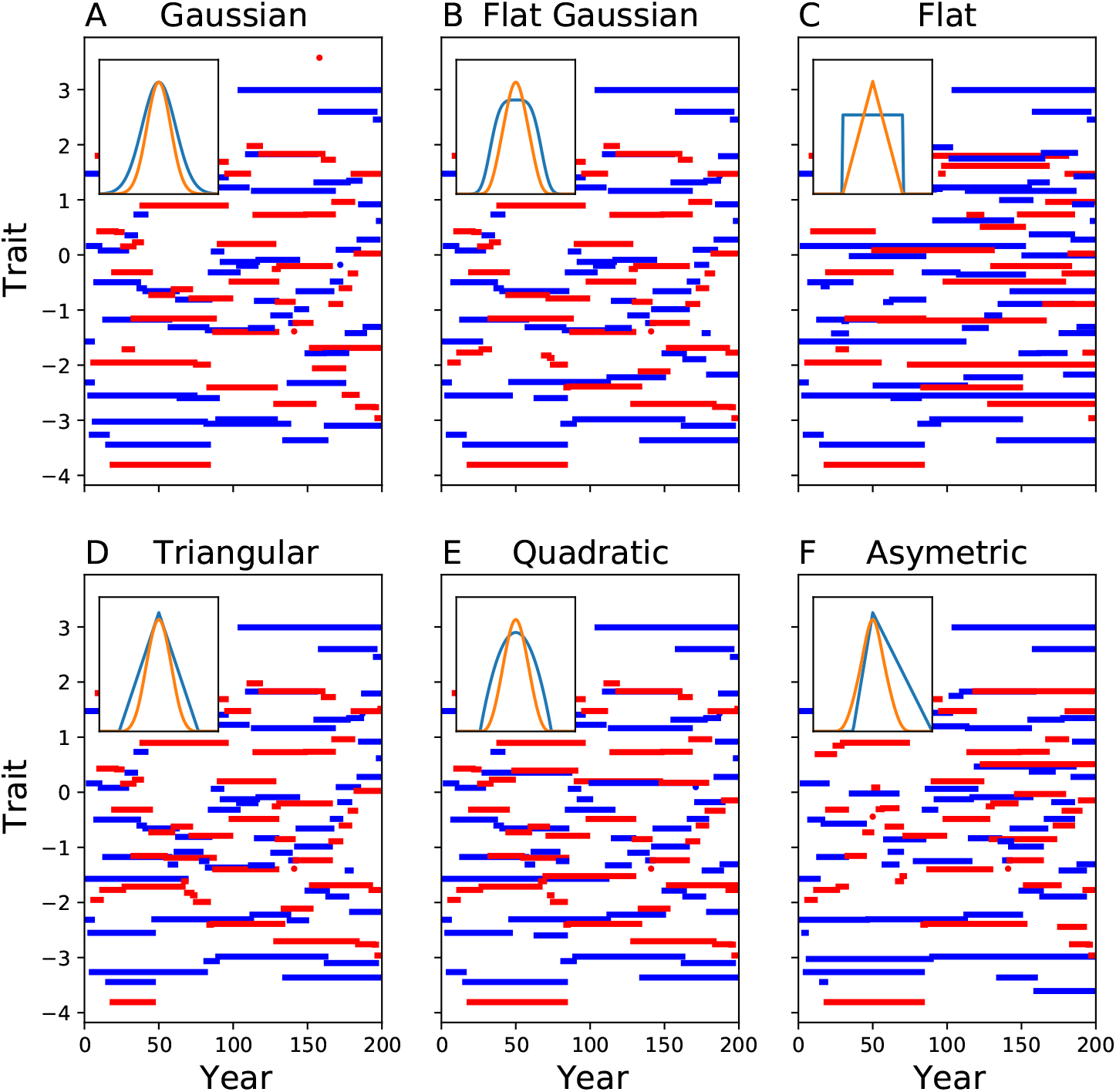
Community dynamics for different resource consumption kernels. For all the different kernels we still observe the continuous community assembly. The inset in each panel shows the resource consumption vector (blue) and the resulting competition kernel for two competing prey species (orange).

**Figure S3:**
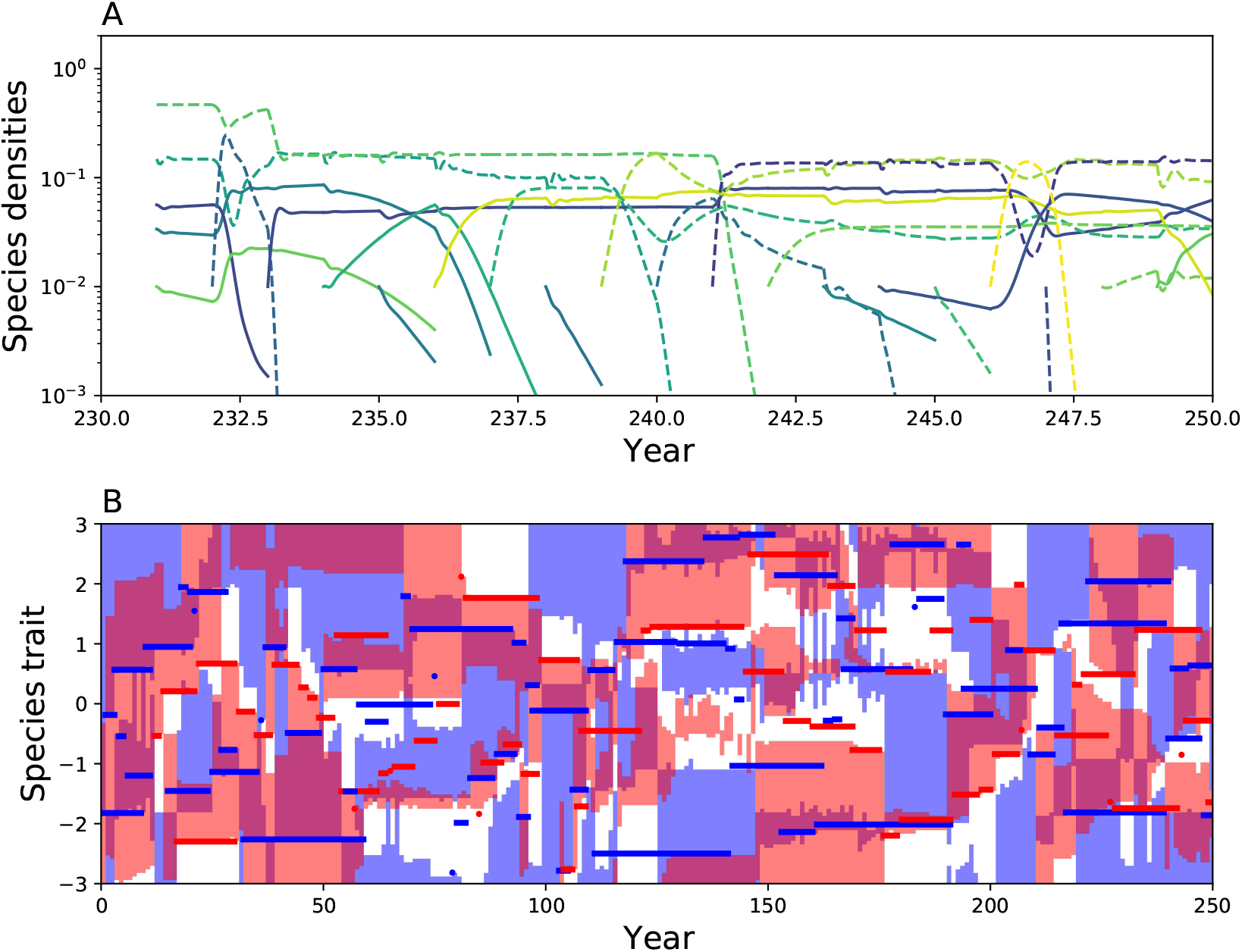
In the main text we did not simulate the community dynamics between invasions, rather we assumed that invasions happen infrequently such that the local community is always at equilibrium when a new species invades. Even if we relax this assumption we still obtain the same the continuous assembly dynamics. A: Densities over time for the last 20 years of the community assembly. Invaders are introduced at density 10^-2^ and go extinct if they fall below 10^-3^ of the total density. B: Species traits of the present species. The shaded areas show where traits of potentially successful invaders.

**Figure S4:**
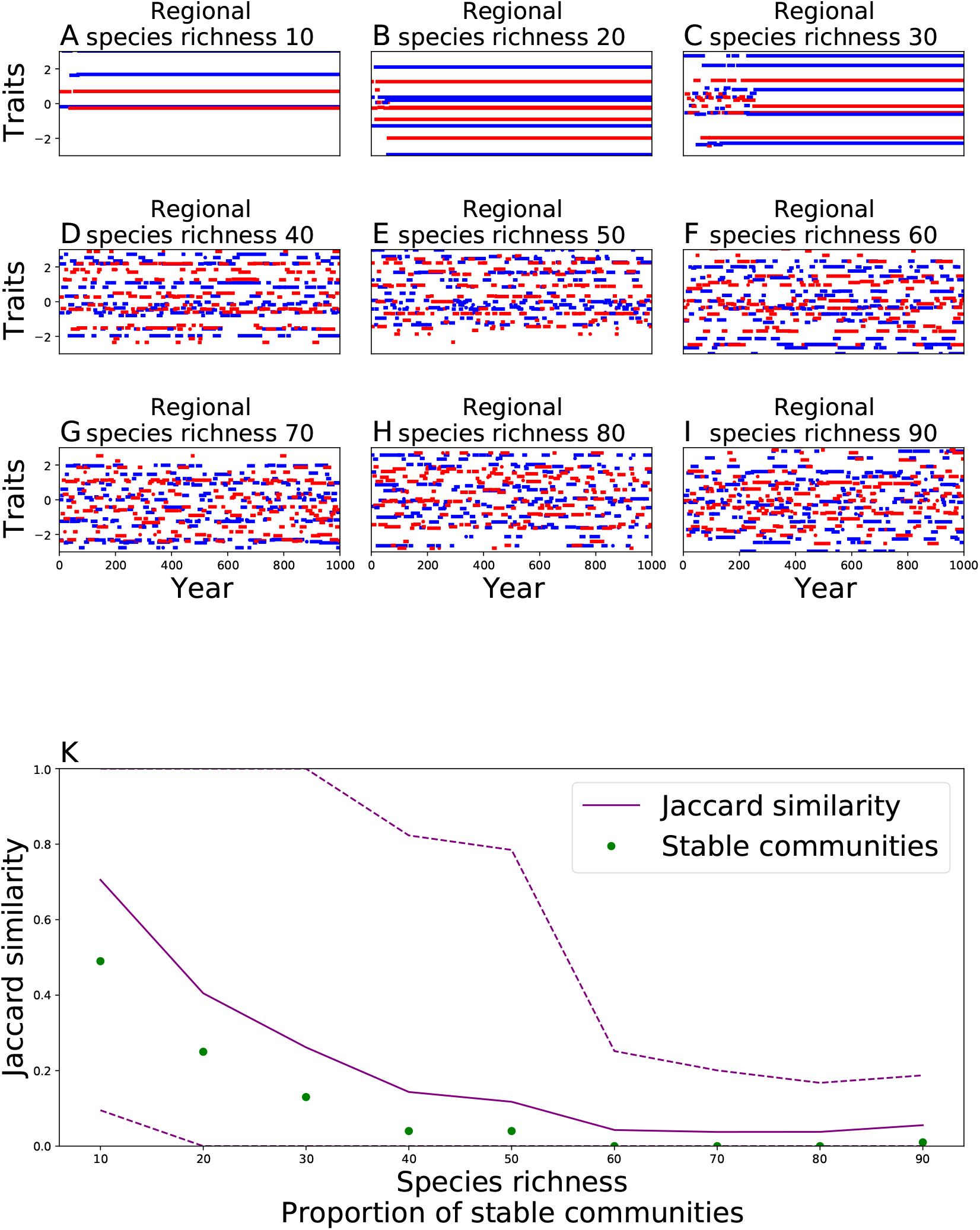
In the main text we assumed an infinite regional species pool. Here we investigate the effects of finite species pools. A-I show sample runs with species richness ranging from 10 to 90 species. Some of these converge towards eventual stability with respect to the regional species pool (A, B and C). The others are also driven by continuous species turnover, although there might be a community composition which is stable in each of these regional species pools, there are 2^n^ possible communities which prohibits a complete search of all possibilities. K: We ran 100 simulations for each species richness. Green dots show the proportion of stable communities, increasing species richness implied lower probability of a stable community. Purple lines show the Jaccard similarity of the end-point with the year 800.

### S4 Additional figures

**Figure S5:**
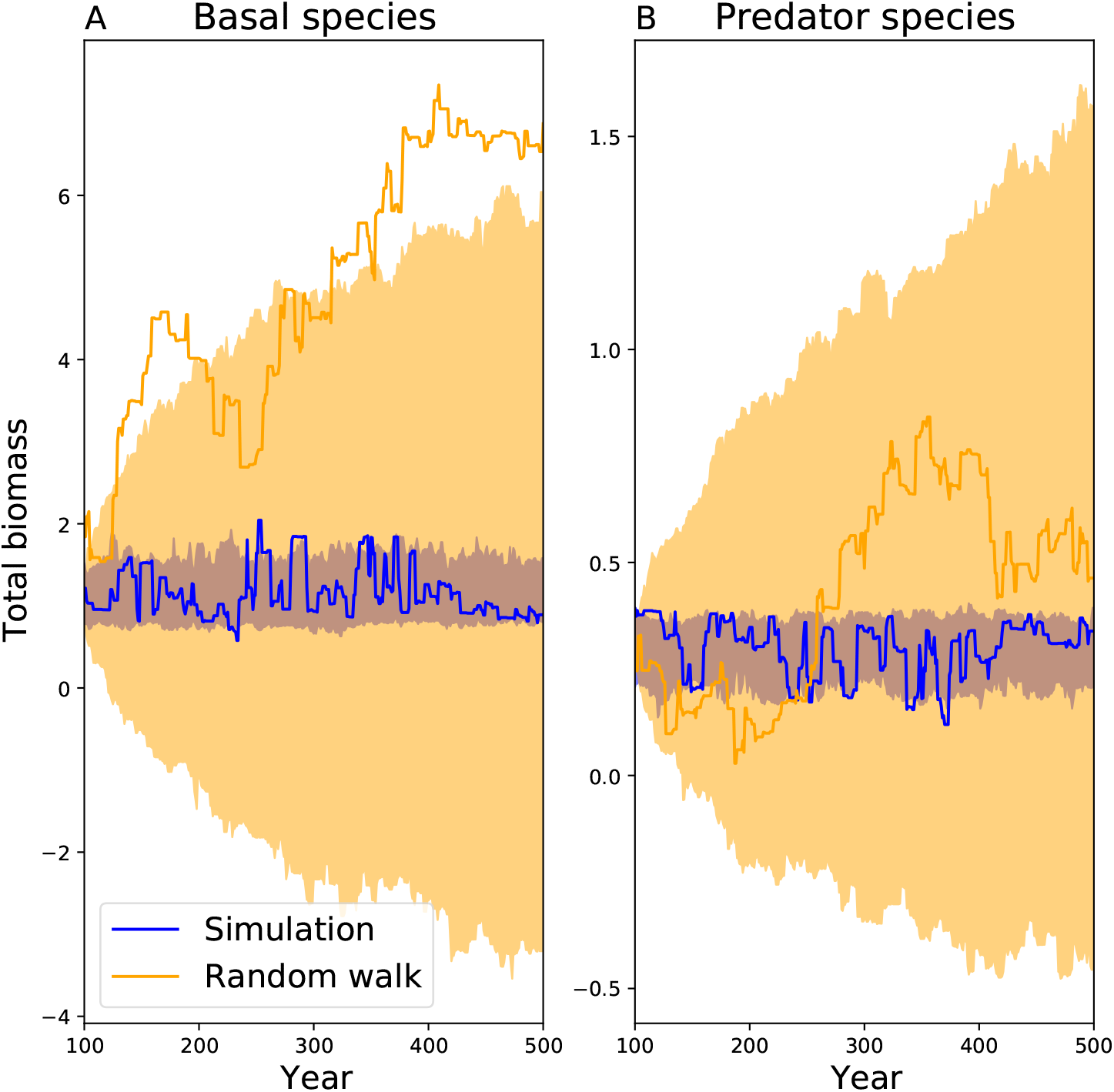
Despite the changes in community composition the total biomass is relatively stable. The blue line shows biomass over time in one specific run, the blue shaded area indicates the 5 and 95% percentile curves of total biomass over multiple runs. We compared this fluctuation in total biomass to a fluctuations in total biomass stemming from a random walk (orange line and shaded area). At each year biomass changes randomly, the changes in biomass are drawn from the actually observed changes in biomass from the community model. As expected, the drift in total biomass in the actual community model is much smaller than the drift in total biomass stemming from the random walk.

**Figure S6:**
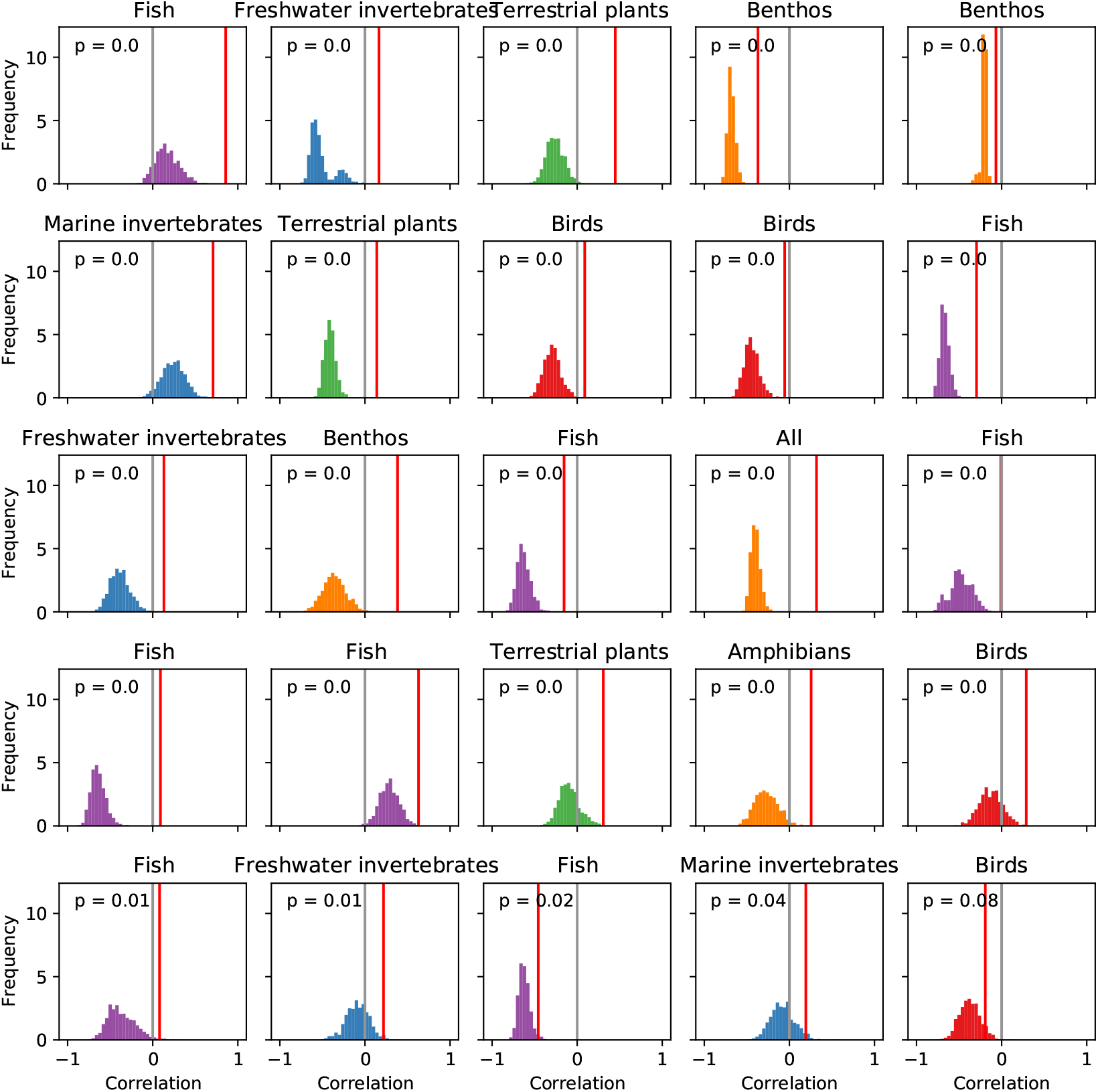
We compare the actual correlation of invasion and extinction in the BioTIME datasets (red vertical line) to the correlation obtained by reshuffling the years 1000 times (histograms). We report the results for the 25 datasets with the lowest p-values (shown in top left corner of each panel).

**Figure S7:**
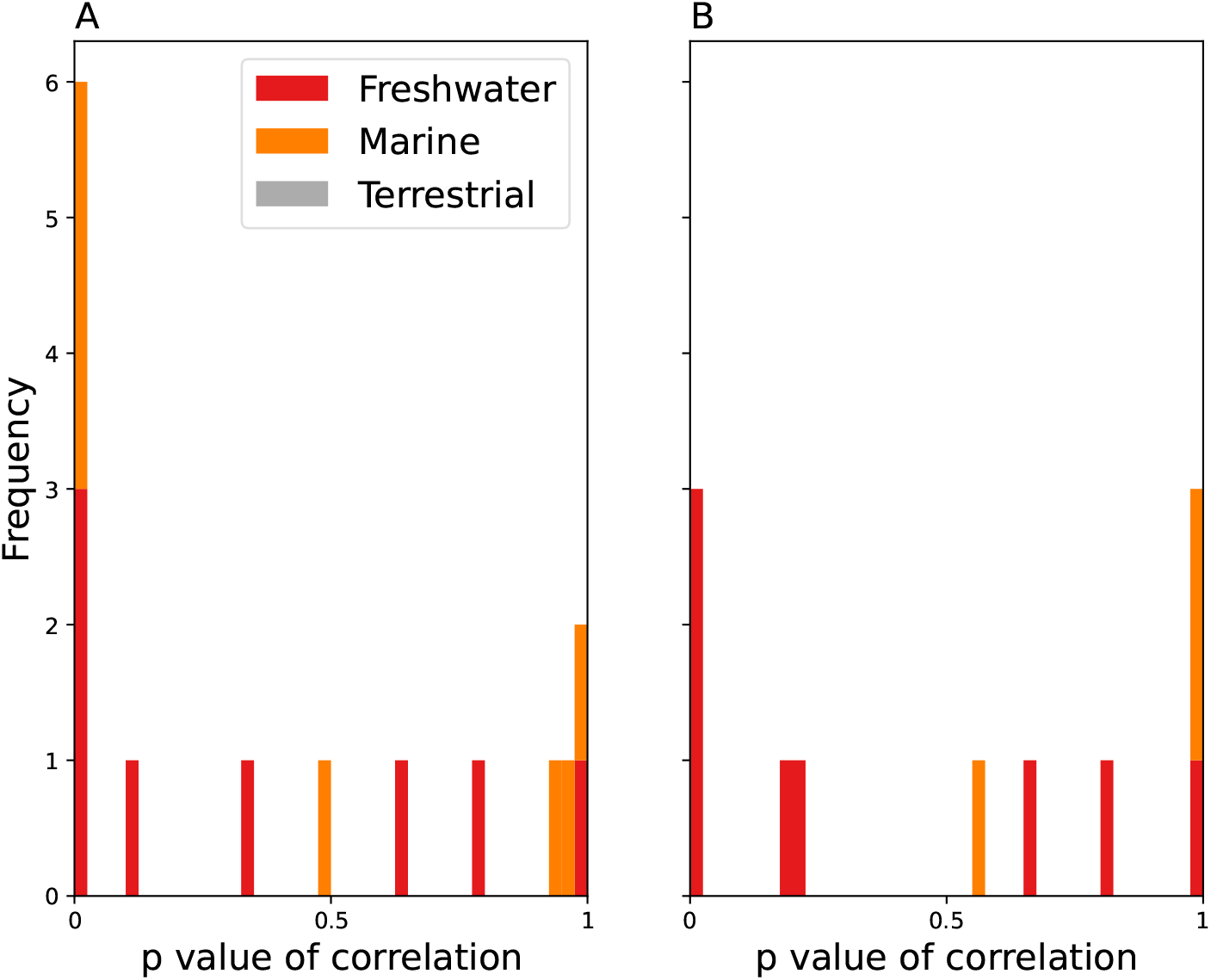
The empirical data from BioTIME contained datasets with very strong fluctuations of species richness (e.g. from over 100 species present to 1 species present within one year). To ensure that our results are not driven by these questionable underlying data we performed two additional tests. Panel A: We have excluded all years in which species richness was below 5 (this threshold was chosen arbitrarily). The total number of datasets remained unchanged by this. Panel B: We have completely excluded all datasets where the maximum species richness is at least four times higher than the minimal species richness, which excluded 18 of the 44 total communities. In both methods we retain the strong correlation of invasions and extinctions.

